# Simple and Highly Specific Targeting of Resident Microglia with Adeno-Associated Virus

**DOI:** 10.1101/2023.12.12.571321

**Authors:** Carolina Serrano, Sergio Cananzi, Tianjin Shen, Lei-Lei Wang, Chun-Li Zhang

## Abstract

Microglia, as the immune cells of the central nervous system (CNS), play dynamic roles in both health and diseased conditions. The ability to genetically target microglia using viruses is crucial for understanding their functions and advancing microglia-based treatments. We here show that resident microglia can be simply and specifically targeted using adeno-associated virus (AAV) vectors containing a 466-bp DNA fragment from the human *IBA1* (*hIBA1*) promoter. This targeting approach is applicable to both resting and reactive microglia. When combining the short *hIBA1* promoter with the target sequence of miR124, up to 95% of transduced cells are identified as microglia. Such a simple and highly specific microglia-targeting strategy may be further optimized for research and therapeutics.

**Significance Statement:** Brain microglia play critical roles in human health and diseases. Genetic manipulation of these cells will offer numerous therapeutic opportunities. However, there is a lack of relevant strategies to target these cells with high specificity since they are traditionally considered to be refractory to virus transduction. Through in vivo screening of many promoters, this study identified a short promoter from the human *IBA1* gene. When incorporated into lentivirus or adeno-associated virus vectors, this promoter proves effective in driving gene expression with high specificity for brain microglia. Such a simple strategy will facilitate specific approaches for microglia-based research.

## INTRODUCTION

Microglia serve as the immune cells of the central nervous system (CNS), maintaining homeostasis under normal conditions. They become activated and play dynamic roles during neuroinflammation, neural damage, or under degenerative conditions (1-6). Depending on the pathological processes, microglia can serve as phagocytes and secrete proinflammatory or anti-inflammatory mediators to regulate degeneration, repair and regeneration in the CNS (7). These behaviors of microglia are tightly regulated by extra- and intra-cellular signaling pathways and transcriptional programs (8, 9). Genetic manipulation of microglia will help understand their function as well as hold promising potential for developing new treatments for CNS diseases.

Viral vectors such as lentivirus and adeno-associated virus (AAV) are clinically relevant tools for genetic manipulation of different cell types (10-13). Even though lentivirus and AAV can be used to transduce most brain cells, microglia seem to be resistant to the transduction using those vectors (13-17). The first analysis using AAV2 showed that microglia could be transduced but gene expression was not detectable, suggesting that microglia could have an intrinsic mechanism to prevent transgene expression or virus degradation (18). The inclusion of microglial specific promoters like F4/80 and CD68 in AAV improved microglia transduction in culture; however, transduction efficiency remained very low in mice (19). Attempts to improve AAV capsids also failed to induce strong and stable microglia transgene expression (20).

In this study, we initially aimed to reprogram microglia for neural regeneration, as we have previously done with resident astrocytes and NG2 glia (21-26). However, multiple attempts failed to efficiently transduce microglia with lentivirus under several routinely used promoters. We therefore conducted in vivo screens of additional promoters that could drive gene expression through lentivirus in resident microglia. We mainly focused on human gene promoters with the expectation that such evolutionarily conserved promoters could eventually be used for therapeutics. Based on the results of lentiviruses, we further revealed that a 466-bp fragment of the human *IBA1* (*hIBA1*) promoter, when employed in AAV vectors, was sufficient to drive microglia-specific gene expression. During the process of this study and supporting our results, it was recently reported that a 1.7-kb mouse *Iba1* promoter was also capable of driving microglia-specific expression through AAVs (27). Notwithstanding, our remarkably short *hIBA1* promoter will allow AAVs to package much larger gene inserts. Although clinically relevant AAV capsids, such as AAV5 and AAV8, can be used to target microglia, our *hIBA1* promoter should be compatible with the recently developed capsid variants for microglia (28, 29).

## RESULTS

### Specific targeting of microglia with lentivirus under the *hIBA1* promoter

We first examined the lentiviral delivery system to target microglia in the adult mouse brain. We screened eight different promoters associated with specific gene expression in microglia. Except for the mouse *mF4/80* promoter, the remaining seven promoters are human origin with an aim of targeting human microglia for clinical therapy. They include the *hCD68, hIBA1, hCX3CR1, hC1QA, hC1QB, hTMEM119*, and *SP* promoters. The reporter *GFP* was put under the control of these promoters, and the lentivirus was packaged with the VSV-G envelope. To include reactive microglia which are critically involved in neurological diseases, we induced ischemic stroke through the MCAO procedure one week before virus injections (Fig. 1A). One week post virus (wpv) delivery, we analyzed the expression of the reporter GFP expression and the microglia-specific marker IBA1 in the injected mouse striatum. GFP^+^IBA1^+^ cells were detected in the striatum with all the promoters tested; however, the highest microglia-specific GFP expression was driven by the *hIBA1* promoter (85.64 ± 0.72 %; Fig. 1B, C). GFP driven by the other promoters was detected in both microglia and cells that morphologically appeared to be neurons and astrocytes. Because the best transduction efficiency for microglia was driven by the *hIBA1* promoter, we decided to focus our subsequent studies on this promoter.

**Figure 1.**
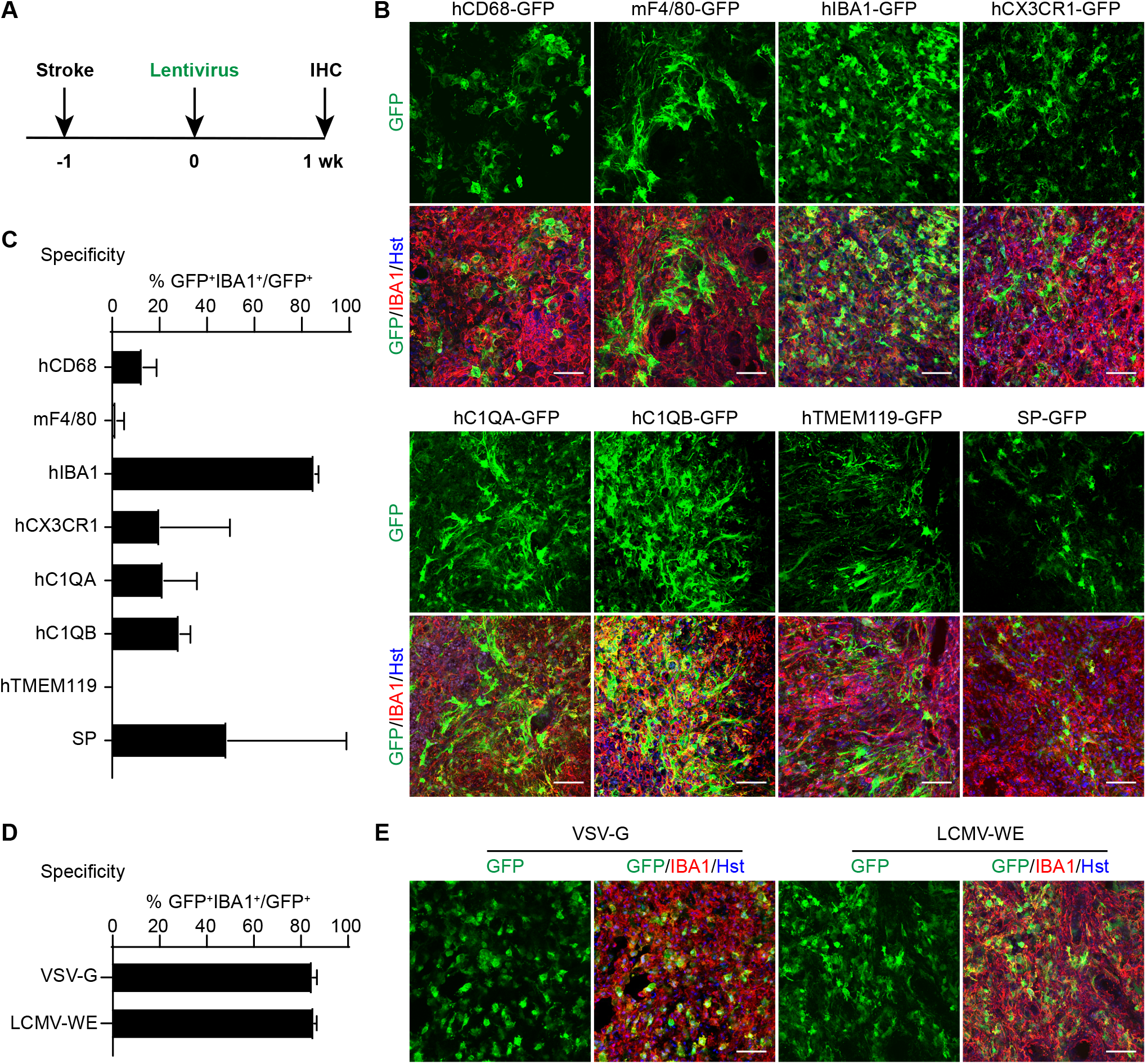
In vivo screens for microglia-targeting lentivirus. (A) Schematic diagram of the experimental procedure. One week post MCAO-induced stroke, entiviruses with various promoter-driven GFP were injected into the striata and examined after another week (wk). (B) Representative confocal images showing marker expression for the indicated lentiviruses. Scale bars, 50 μm. (C) Quantifications showing high microglia-specificity of GFP expression in mice injected with the *enti-hIBA1-GFP* (mean ± SEM; n=3 mice per group). (D) Quantifications showing comparable specificity of lentivirus packaged with either VSV-G or LCMV-WE envelope (mean ± SEM; n=3 mice per group). (E) Representative confocal images showing marker expression for the indicated lentiviruses. Scale bars, 50 μm.

### Microglia-specific expression is not affected by the lentiviral envelopes

Cell type-specificity of lentiviruses was reported to be influenced by the viral envelopes (30). To test this possibility, we compared the *hIBA1-GFP* lentivirus packaged with either the VSV-G or the LCMV-WE envelope. When GFP^+^IBA1^+^ cells were examined at 1 wpv, both enveloped lentiviruses showed high targeting specificity for microglia in the injected striatum (84.96 ± 1.04% for VSV-G and 85.64 ± 0.75 % for LCMV-WE; Fig. 1D, E).

### Specific targeting of microglia with scAAVs under the *hIBA1* promoter

Lentiviral and AAV vectors are widely applied for gene expression *in vivo*. Even though both systems allow stable transgene expression, AAVs are more broadly employed for *in vivo* gene therapy due to their non-integration nature and minimal inflammatory response (11, 12, 31, 32). We first examined the self-complementary AAV (scAAV) since it was reported to drive stronger gene expression than the single strand AAV (ssAAV) (33). *scAAV-hIBA1-GFP* was then packaged with 8 different serotypes, including AAV1, AAV2, AAV5, AAV6, AAV6m, AAV8, AAV9 and PHP.eB. They were injected into the striatum one week post L-NIO-induced stroke (which produces more constant and confined infarcts when compared to the MCAO procedure) and examined 7 days later (Fig. 2A). All these scAAVs produced a robust expression of GFP in IBA1^+^ cells but with varying degrees of specificity and efficiency (Fig. 2B-D). Quantification of GFP^+^IBA1^+^ cells showed that both scAAV5 and scAAV8 had the highest degrees of specificity (93.68 ± 1.14% and 77.42 ± 4.97%, respectively; Fig. 2C) and efficiency (69.7 ± 7.23% and 82.11 ± 0.8%, respectively; Fig. 2D) for microglia.

**Figure 2.**
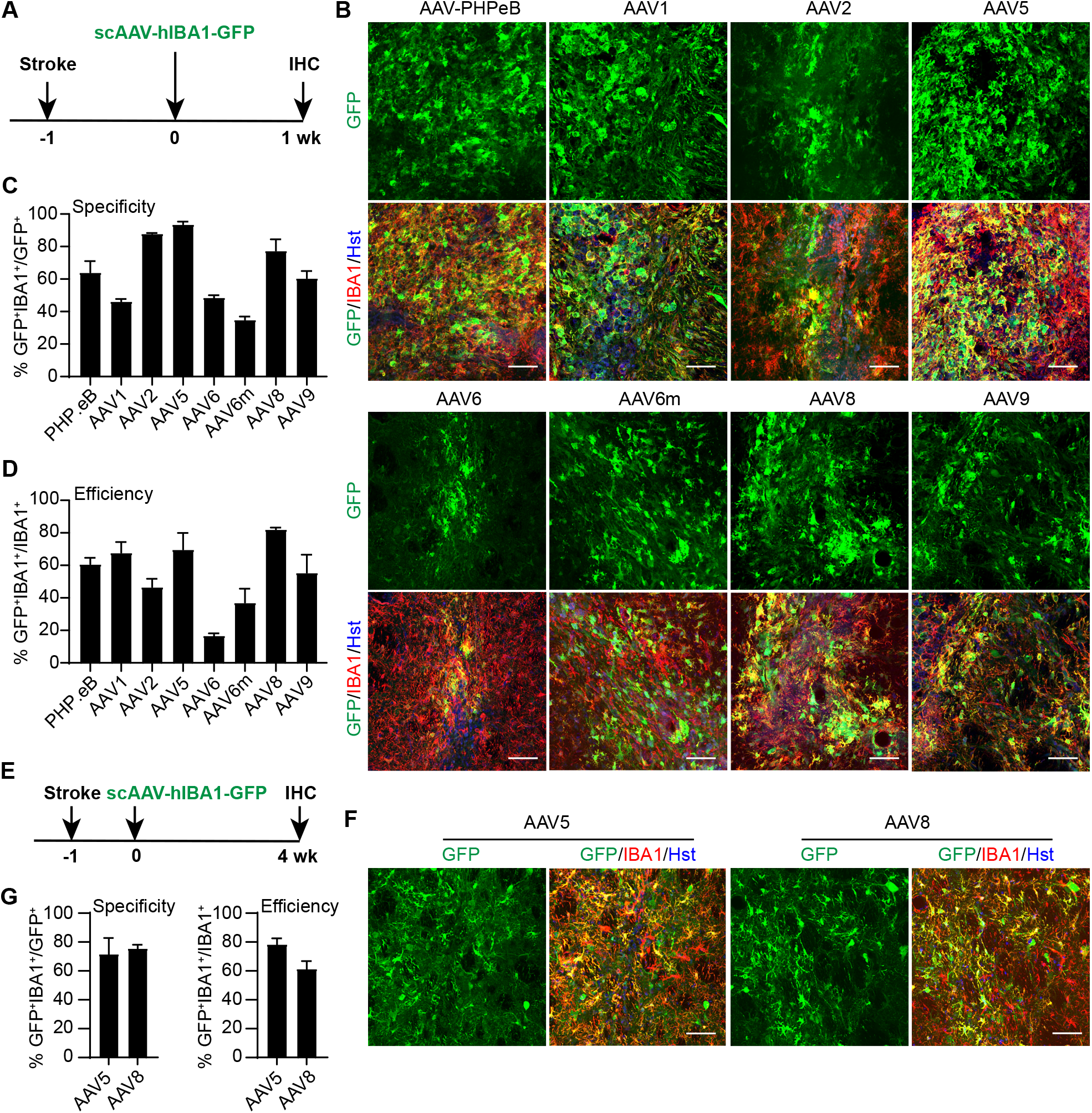
In vivo screens for microglia-targeting AAVs. (A) Schematic diagram of the experimental procedure. One week post L-NIO-induced stroke, scAAVs with various serotypes were injected into the striata and examined after another week (wk). (B)Representative confocal images showing marker expression for the indicated scAAVs. Scale bars, 50 μm. (C) Quantifications showing microglia-specificity of GFP expression for the indicated scAAVs (mean ± SEM; n=3 mice per group). (D) Quantifications showing microglia transduction efficiency for the indicated scAAVs (mean ± SEM; n=3 mice per group). (E) Schematic diagram of the experimental procedure. One week post L-NIO-induced stroke, scAAV5 or scAAV8 was injected into the striata and examined after another 4 week (wk). (F) Representative confocal images showing marker expression for the indicated scAAVs. Scale bars, 50 (G) Quantifications showing the specificity and transduction efficiency for microglia (mean ± SEM; n=3 μm.

To determine whether transgene expression could be maintained for longer time in vivo, we injected the virus and conducted analysis at 4 wpv (Fig. 2E). Both scAAV5 and scAAV8 could mediate robust GFP expression in microglia at this longer delay after virus injection (Fig. 2F-G). Cell type-specificity and transduction efficiency were comparable to what were observed at 1 wpv.

### Identification of the minimal *hIBA1* promoter

The AAV vector has a limited packaging capacity, with about ∼4.5 kb for ssAAVs and an even smaller size for scAAV (34, 35). A minimal promoter will help increase the size of the transgene that could be packaged. Our original *hIBA1* promoter is about 760 bp in length, which is already relatively small. To determine whether it could be further shortened, we made a series of truncations starting from the 5’ end and subcloned them into the ssAAV vector. They were then packaged with serotype 5 and injected into the striatum of adult mouse with or without L-NIO-induced stroke (Fig. 3A-B). When examined at 1 wpv, the 466-bp *hIBA1a* promoter showed a similar expression pattern as the original *hIBA1* promoter, with high microglia specificity and transduction efficiency (70.51 ± 5.12% and 82.26 ± 2.62%, respectively; Fig. 3C-E). Of note, the microglia-specificity was comparable in brains with or without the stroke (70.51 ± 5.15% for sham and 91.93 ± 1.98% for stroke; Fig. 3D), although the transduction efficiency tended to be slightly lower than the non-stroke condition (Fig. 3E). The other shorter promoters, especially *hIBA1c* and *hIBA1d*, performed worse. These results suggest that the 466-bp *hIBA1a* promoter could be used for subsequent experiments. When examined at 4 wpv, however, all these viruses showed a lower microglia-specificity and transduction efficiency (Fig. S1), indicating a time-dependent effect for the ssAAV virus.

**Figure 3.**
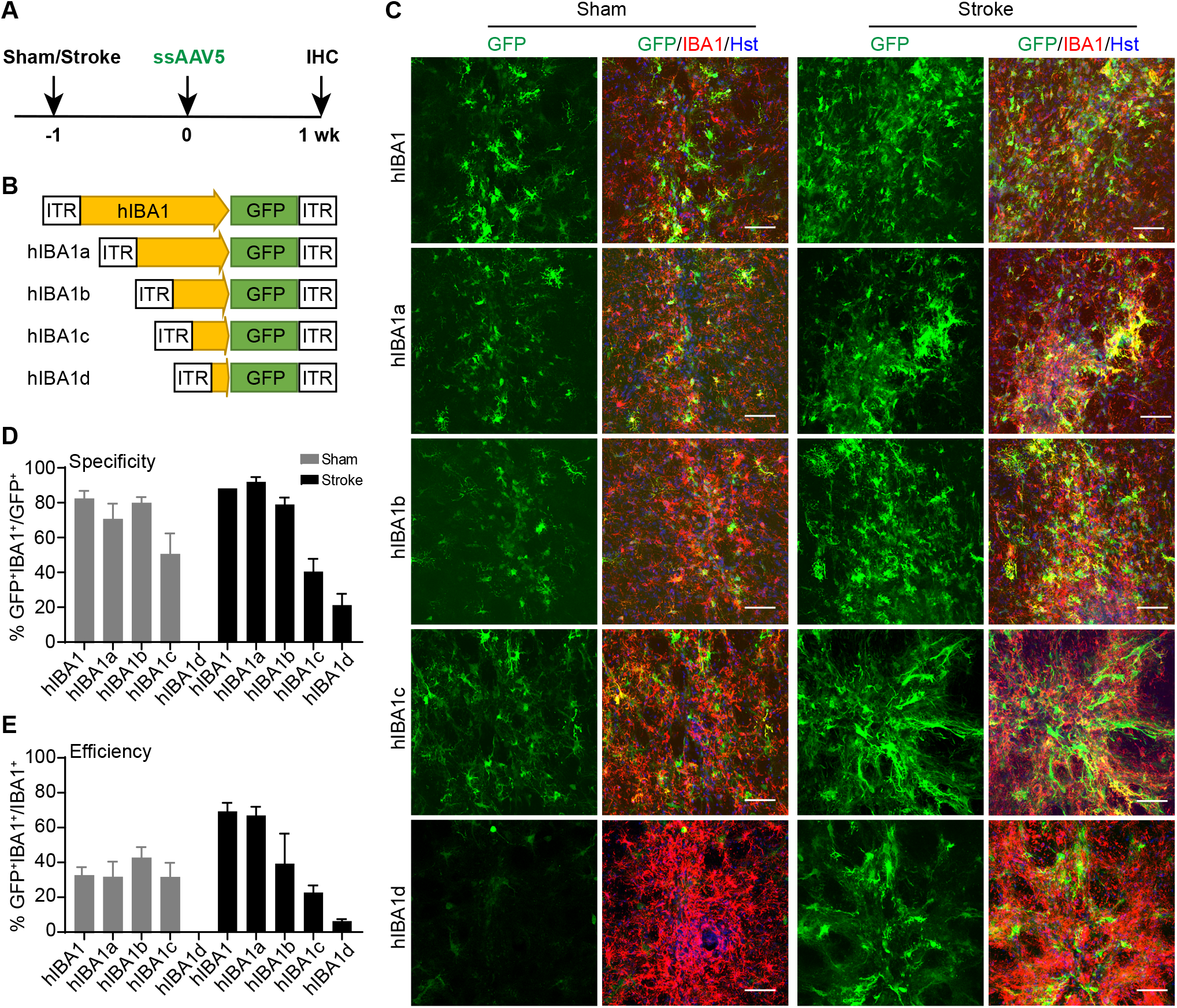
In vivo screens for the minimal microglia-targeting promoter. (A) Schematic diagram of the experimental procedure. ssAAV5 viruses with different promoters were njected into sham mice or mice with L-NIO-induced stroke. Brains were analyzed one week later. (B) Diagram of the examined ssAAVs with different lengths of *hIBA1* promoter. (C) Representative confocal images showing marker expression for the indicated ssAAV5s. Scale bars, 50 μm. (D) Quantifications showing microglia-specificity of GFP expression for the indicated ssAAV5s (mean ± SEM; n=3 mice per group). (E) Quantifications showing microglia transduction efficiency for the indicated ssAAV5s (mean ± SEM; n=3 mice per group).

### High microglia-specificity of scAAV under the *hIBA1a* promoter

Since the above ssAAVs had dramatically reduced microglia specificity and transduction efficiency at 4 wpv, we then evaluated whether packaging the virus as scAAV would improve the outcome. For this purpose, we prepared the *scAAV5-hIBA1a-GFP* virus, injected it into the striatum of adult mouse with or without L-NIO-induced stroke, and examined gene expression at both 1 and 4 wpv (Fig. 4A). We found strong GFP expression in microglia at 1 wpv, in both non-injured and injured brains (Fig. 4B). The efficiency and specificity for microglia transduction was not only maintained but also slightly increased at 4 wpv, reaching nearly 80% under the stroke condition (87.62 ± 1.16% for specificity and 88.74 ± 1.12% for efficiency; Fig. 4B-D). These results suggest that the *hIBA1a* promoter could drive highly microglia-specific expression when packaged into scAAV.

**Figure 4.**
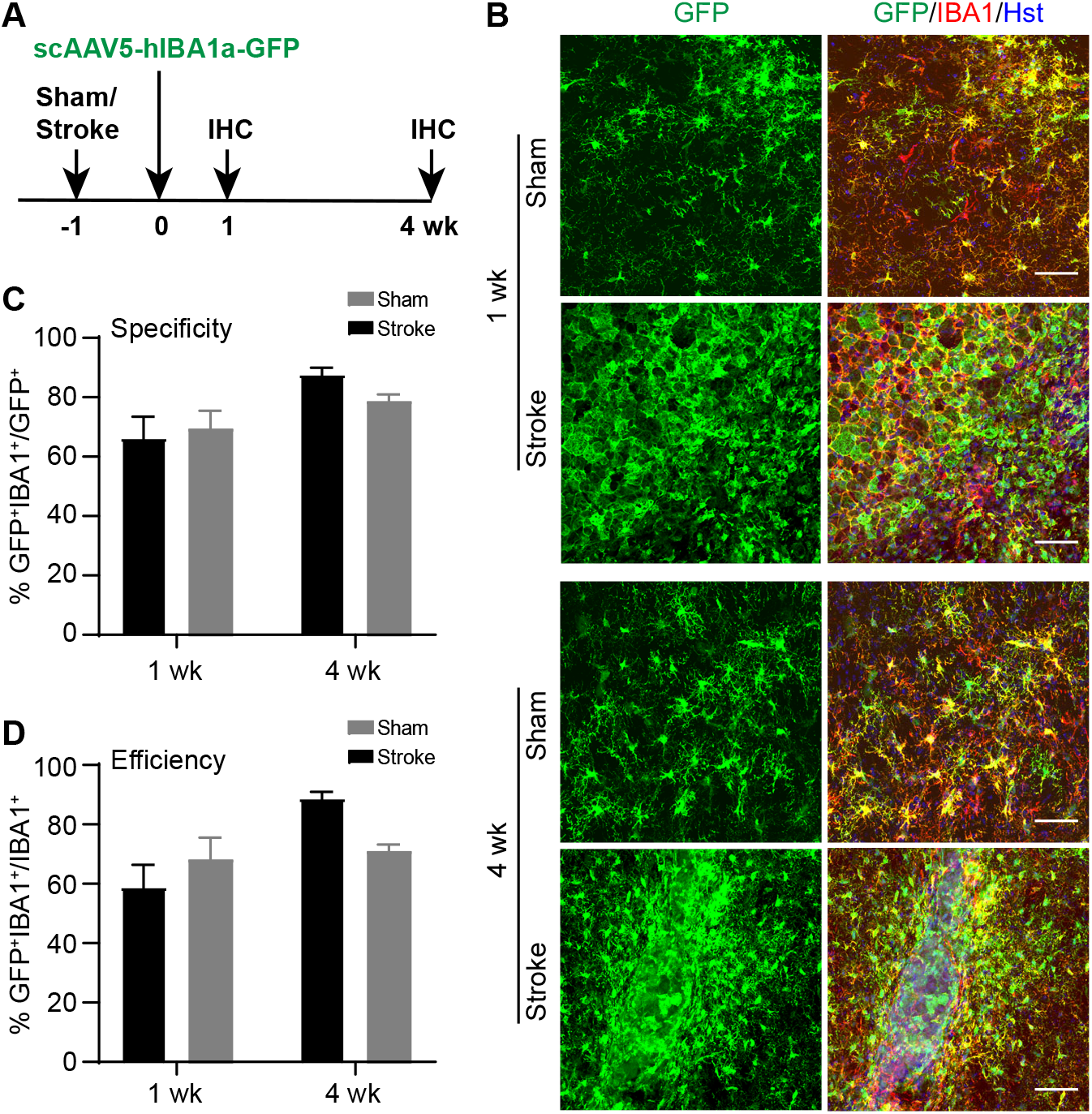
Increased long-term specificity and efficiency for the minimal *hIBA1a* promoter when packaged in scAAV. (A) Schematic diagram of the experimental procedure. scAAV5 virus with the *hIBA1a* promoter was njected into sham mice or mice with L-NIO-induced stroke. Brains were analyzed 1 week or 4 weeks ater. (B) Representative confocal images showing marker expression for the indicated conditions. Scale bars, 0 μm. (C) Quantifications showing microglia-specificity of GFP expression for the indicated conditions (mean ± SEM; n=3 mice per group). (E) Quantifications showing microglia transduction efficiency for the indicated conditions (mean ± SEM; n=3 mice per group).

### miR124T confers high specificity of ssAAV under the *hIBA1a* promoter

Although scAAV can improve the stability for microglia transduction, its limited packaging size could be a problem for larger genes. To maintain microglia specificity but also keep higher packaging capacity, we redesigned a new ssAAV vector under the *hIBA1a* promoter. Since the vast majority of GFP^+^ non-microglia cells were neurons, we inserted after the transgene a synthetic sequence containing 4 copies of the targeting sequence of miR124 (Fig. 5A, miR124T). miR124 is a microRNA that is highly enriched in neurons (36). The insertion of miR124T after the transgene is expected to cause silencing of the transgene in neurons but not in other cells (37). We packaged the *ssAAV5-hIBA1a-GFP-miR124T* virus and injected it into the striatum of mouse without prior injury (Fig. 5A). When examined at 4 wpv, we observed robust and highly specific GFP expression in IBA1^+^ microglia (Fig. 5B). Quantification showed that 94.78 ± 0.59% of GFP^+^ cells were also IBA1^+^ (Fig. 5C), indicating a remarkably high microglia-specificity. The remaining few GFP^+^ but IBA1^-^ cells mainly exhibited the morphology of astrocytes or neurons. Transduction efficiency surrounding the injected brain area was also high, reaching 85.83 ± 1.79% (Fig. 5D).

**Figure 5.**
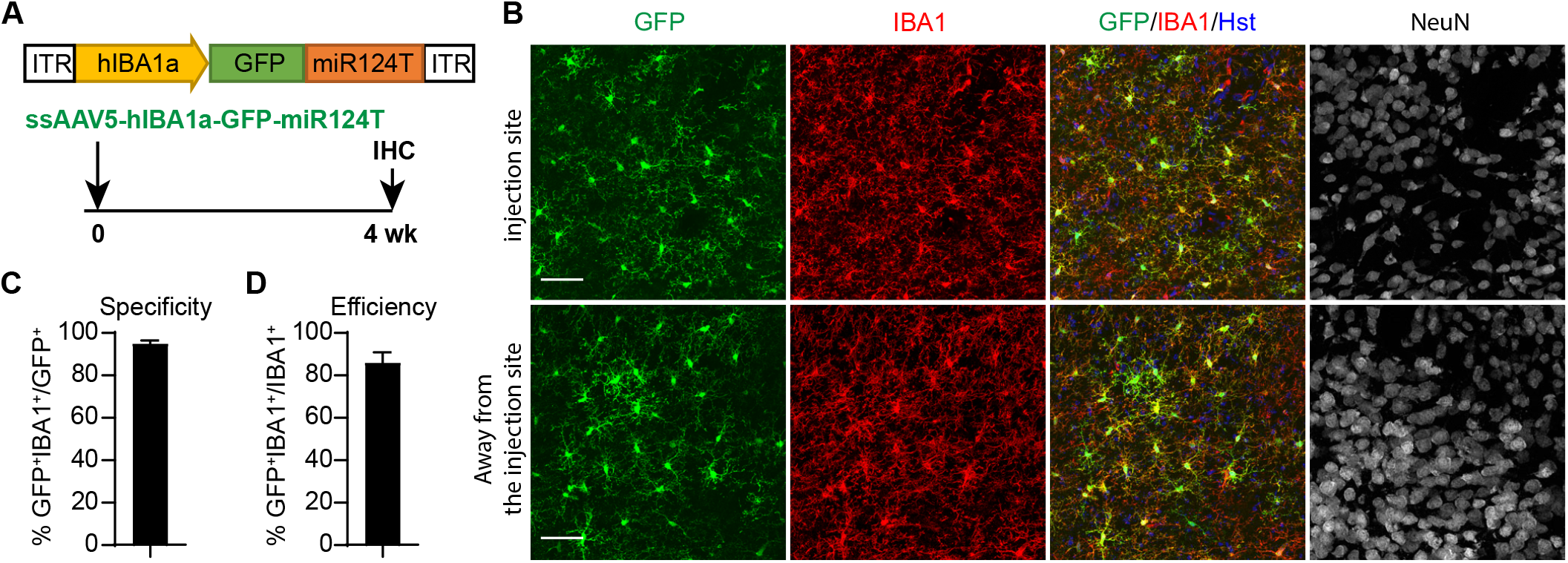
miR124T confers high specificity of ssAAV under the *hIBA1a* promoter. (A) Schematic diagram of the experimental procedure. The *miR124T* sequence was inserted into the 3’ end of the GFP gene. ssAAV5 virus was then injected into the striatum of adult wildtype mouse and analyzed 4 weeks later. (B) Representative confocal images showing marker expression in the injection area and an area away from the injection site. Neurons are marked by the NeuN staining. Scale bars, 50 μm. (C) Quantifications showing high microglia-specificity of GFP expression (mean ± SEM; n=3 mice per group). (D) Quantifications showing high microglia transduction efficiency (mean ± SEM; n=3 mice per group).

## DISCUSSION

Our results show that brain microglia can be specifically targeted with AAVs containing a 466-bp *hIBA1* promoter (*hIBA1a*). The microglia specificity is further improved when the target sequence of miR124 (miR124T) is included in the AAV vector to inhibit transgene expression in neurons. The *hIBA1a* promoter and miR124T could be similarly integrated into lentiviral vectors for lentivirus-mediated targeting of resident microglia.

Although it is well-known that viral tropism is determined by surface proteins in the viral envelope or capsid, cell type-specificity of transgene expression is also controlled by the gene promoters (12, 13, 38, 39). In lentiviral vectors, our in vivo screens showed that the GFP reporter driven by the *hIBA1* promoter exhibited the highest microglia-specificity when compared to any other tested promoters, including the synthetic promoter (*SP*), *hCD68, mF4/80, hCX3CR1, hC1QA*, and *hC1QB*. Future experiments may be needed to tease out the regulatory genomic elements that drive microglia-restricted expression by these latter genes. Of note, microglia-specificity of lentiviral vectors with the *hIBA1* promoter was not majorly affected by the viral envelope, as we found a comparable specificity for lentivirus packaged with either the VSV-G or the LCMV-WE envelope. Such a result further highlights the importance of the specific promoter for gene expression in microglia.

Because of its low toxicity and minimal induction of host immune responses, AAVs are broadly employed for gene expression *in vivo (11, 32)*. Among the many natural capsids of AAVs, AAV1, AAV2, AAV5, AAV6, AAV8, and AAV9 are the most frequently used for research and therapeutics. Our results showed that AAV vector containing the *hIBA1* promoter, when packaged with any of these above capsids, could drive transgene expression in microglia. Nonetheless, AAV5 and AAV8 gave rise to a comparably high transduction specificity and efficiency for microglia, while the previously reported AAV6 mutant capsid (AAV6m) (20) performed the worst. The non-microglia cells for these above tested capsids are mainly neurons and astrocytes, which seem to be the default cell types for majority AAVs even with glia-specific promoter (15, 40). To reduce non-specific transgene expression in neurons, we inserted into the AAV vector with 4 copies of the target sequence of miR124, a microRNA highly enriched in neurons (36). Such a strategy greatly improved targeting specificity and long-term expression through AAVs in brain microglia; and it could be similarly employed in the lentiviral vectors.

A major limitation of AAVs is their relatively small packaging capacity for foreign DNAs, with roughly 4.5 kb for ssAAVs (34) and an even smaller size for scAAVs (35). Our systematic truncation analysis showed that a 466-bp genomic fragment of the *hIBA1* promoter is still sufficient to drive microglia-specific gene expression, therefore permitting insertion of larger genes in both ssAAV and scAAV vectors. Our result is in sharp contrast to a recent report, showing a 1.7-kb fragment of the mouse *Iba1* promoter is needed for expression in microglia (27). Additionally, our short *hIBA1* promoter is expected to work in human microglia; therefore, it should have broader applications than the much longer mouse *Iba1* promoter.

In conclusion, our study identified a simple method to specifically target resident microglia with AAVs or lentiviruses. It could be further facilitated with mutant AAV capsids for microglia (28, 29). Our method should be a valuable contribution to microglia-based research and therapeutics.

## MATERIALS AND METHODS

### Animals

Wildtype C57/BL6J mice were purchased from the Jackson Laboratory. Adult male and female mice at 2-3 months of age were used unless otherwise stated. All mice were housed under a 12 h light/dark cycle and had *ad libitum* access to food and water in the animal facility. Experimental animals were randomized, and the experimenters were not blinded to the allocation of animals during experiments and outcome assessment. All experimental procedures and protocols were approved by the Institutional Animal Care and Use Committee at University of Texas Southwestern.

### Stroke

Strokes were induced by either middle cerebral artery occlusion (MCAO) or L-NIO injection. For MCAO, mice were anesthetized with isoflurane. The left common and external carotid arteries were exposed and ligated. Occlusion of the middle cerebral artery (MCA) was accomplished through the insertion of a filament from the basal part of the external carotid artery and advancing it in the internal carotid artery toward the location of MCA branching from the circle of Willis. The occlusion lasted 20 min and reperfusion was initiated by removing the suture from the internal carotid artery. Once the suture was removed, the internal carotid artery was ligated. L-NIO induced ischemia was selected for the following experiments because it produced constant and confined infarcts in the striatum. For L-NIO induced ischemia, a craniotomy was performed overlying the injection sites of the cortex. A Hamilton syringe was filled with L-NIO (27 *μ*g/*μ*L in sterile physiological saline; Calbiochem), secured onto the stereotaxic arm, and connected to a pressure pump. Three injections (each of 0.3 *μ*L of L-NIO solution) were made in the following coordinates: anterior-posterior (AP) +1 mm, medio-lateral (ML) +2 mm, dorsoventral from the skull (DV) −2 mm; AP +1 mm, ML +2 mm, DV −2.5 mm; and AP +1 mm, ML +2 mm, DV −3 mm. Injections were made at a rate of 3 *μ*L/min, targeting the striatum. Localized vasoconstriction leads to focal ischemia in the striatum. In Sham experiments, the needle was placed at the same coordinates as the stroke experiments, but L-NIO was not injected in the striatum.

### Lentiviral vectors and virus preparation

The lentiviral vector SP–GFP was generated by sub-cloning the macrophage synthetic promoter (SP, ∼400 bp, which harbors multiple myeloid/macrophage cis elements) into the CS–CDF–CG–PRE vector (41). This SP-GFP vector was used to construct other lentiviral vectors. The human *CD68* promoter and enhancer (*hCD68*, ∼800 bp) was subcloned from *pcDNA3-hCD68prm* (Addgene #34837). The mouse *F4/80* promoter (*mF4/80*, ∼670 bp) was PCR-amplified from mouse genomic DNA. The human *IBA1* promoter (*hIBA1*, ∼760 bp), human *TMEM119* promoter (*hTMEM119*, ∼570 bp), human *CX3CR1* promoter (*hCX3CR1*, ∼700 bp), human *C1QA* promoter (*hC1QA*, ∼560 bp), and human *C1QB* promoter (*hC1QB*, ∼550 bp) were PCR-amplified from human genomic DNA. All vectors were verified through restriction enzyme digestions and DNA sequencing. Replication-deficient virus was produced in HEK293T cells by transient transfection with lentiviral vectors and packaging plasmids (pMDL, pREV, and VSV-G or pHCMV-LCMV-WE envelopes). Lentivirus was collected by PEG precipitation and tittered as described previously (21).

### AAV vectors and virus production

The single strand AAV (ssAAV) vectors were based on the *AAV-hGFAP-GFP* vector (40) by replacing the *hGFAP* promoter with the *hIBA1* promoter. The self-complementary AAV (scAAV) vectors were based on the *scAAV-CAG-GFP* vector (Addgene #83279) by replacing the *CAG* promoter with the *hIBA1* promoter. All vectors were verified through restriction enzyme digestions and DNA sequencing. AAV viruses were packaged with *pAd-deltaF6* (Addgene #112867) and one of the following helper plasmids: *pAAV2/1* (Addgene #112862), *pAAV2/2* (Addgene #104963), *pAAV2/5* (Addgene #104964), *pAAV2/6* (Addgene #110660), *pAAV2/6m* (20), *pAAV2/8* (Addgene #112864), *pAAV2/9* (Addgene #112865), or *pUCmini-iCAP-PHP*.*eB* (Addgene #103005). Briefly, HEK293T cells were transfected with the packaging plasmids and a vector plasmid. Three days later, virus was collected from the cell lysates and culture media. Virus was purified through iodixanol gradient ultracentrifugation, washed with PBS, and concentrated with 100K PES concentrator (Pierce™, Thermo Scientific). Viral titers were determined by quantitative PCR with ITR primers (forward: 5-GGAACCCCTAGTGATGGAGTT-3; reverse: 5-CGGCCTCAGTGAGCGA-3).

### Stereotactic brain injections

Viruses were stereotactically injected into the striata of adult mice as described previously (24). Briefly, mice were placed on a stereotactic frame (Kopf, USA) under isoflurane anesthesia. A vertical skin incision exposed the Bregma on the skull used for guiding the location of the Burr hole. Injection coordinates were as follows: AP +1.0 mm, ML -2.0 mm, and DV -3.0 mm. 2 *μ*L of lentivirus with an original titer of ∼2e9 cfu/mL or 2 *μ*L of AAV with an original titer of 0.5-8e13 GC/mL was injected using a 10-*μ*L Hamilton syringe and a 33-gauge beveled tip metal needle. Virus was injected at a rate of 0.5 *μ*L/min until the total volume was delivered. The needle was left in place for 5 minutes before it was slowly removed from the brain.

### Immunohistochemistry and quantification

Mice were sacrificed and fixed by intracardial perfusion with 4% paraformaldehyde in PBS. Brains were dissected out, post-fixed overnight and cryoprotected with 30% sucrose at 4°C for 48 hrs. Coronal brain sections were sectioned on a sliding microtome set at 40 *μ*m thickness. The following primary antibodies were used: GFP (GFP-1020, chicken, 1:1000, Aves Labs), GFAP (ab4674, chiken, 1:500, Abcam), IBA1 (019-19741, rabbit, 1:1000, Waco), and NeuN (266 004, guinea pig, 1:500, Cedarlane-synaptic system). Alexa Fluor 488-, 594- or 647-conjugated corresponding secondary antibodies from Jackson ImmunoResearch were used for indirect fluorescence. Nuclei were counterstained with Hoechst 33342 (Hst). Images were captured and examined by a Zeiss LSM 700 confocal microscope. Three to five random confocal images with marker-positive cells around the injection side were analyzed for each animal. Microglia-specificity was calculated as % of GFP^+^IBA1^+^/IBA1^+^ cells, whereas viral transduction efficiency was calculated as % of GFP^+^IBA1^+^/IBA1^+^ cells.

## DECLARATIONS

### Availability of data and material

All data generated or analyzed during this study are included in the article or its supplementary information files.

### Authors’ contributions

C.S., S.C. and C.-L.Z. conceived and designed the experiments. C.S. and S.C. performed research, created figures, and drafted the original manuscript. T.J. and L.-L.W. provided essential technical support. C.-L.Z revised the manuscript. All authors reviewed and approved the final manuscript.

## Acknowledgments

We thank members of the C.-L.Z. laboratory for helpful discussions, reagents and technical assistance, Yuhua Zou for maintaining mouse colonies, and Paramita Chakrabarty for providing AAV6 mutant capsid.

