## Supplemental Figures for "Simple and Highly Specific Targeting of Resident Microglia with Adeno-Associated Virus"

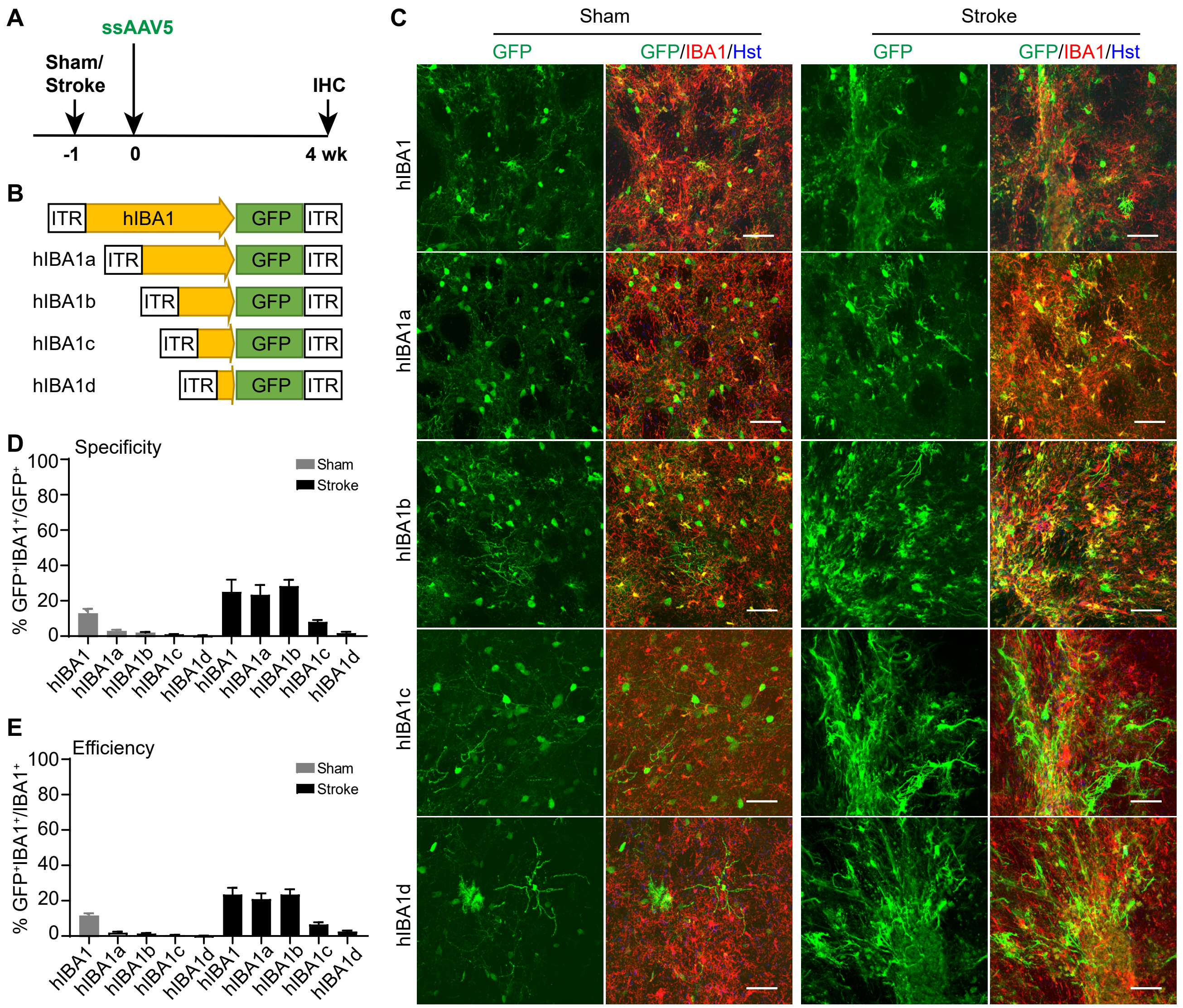


**Figure S1. Reduced transduction specificity and efficiency at a longer delay post ssAAV injection.** (A) Schematic diagram of the experimental procedure. ssAAV5 viruses with different promoters were injected into sham mice or mice with L-NIO-induced stroke. Brains were analyzed 4 weeks later.

(B) Diagram of the examined ssAAVs with different lengths of *hIBA1* promoter.

(C) Representative confocal images showing marker expression for the indicated ssAAVs. Scale bars, 50 µm.

(D) Quantifications showing microglia-specificity of GFP expression for the indicated ssAAVs (mean ± SEM; n=3 mice per group).

(E) Quantifications showing microglia transduction efficiency for the indicated ssAAVs (mean ± SEM; n=3 mice per group).
